# Determining perception thresholds of young adults to small continuous moving platform perturbations

**DOI:** 10.1101/2025.08.21.671548

**Authors:** Kimia Mahdaviani, Luc Tremblay, Alison Novak, Avril Mansfield

## Abstract

Detecting external disturbances is vital for maintaining balance, as corrective actions are initiated to prevent falls. Quantifying people’s ability to perceive such disturbances improves our understanding of how balance is maintained. This study aims to: 1) quantify healthy young adults’ ability to perceive external perturbations while balancing on a stabilometer, and 2) understand the relationship between balance performance and perturbation magnitude relative to participants’ perception threshold. Participants (n=22; 20–35 years) completed a multiple staircase protocol. While standing on a stabilometer mounted on a moving platform, they attempted to keep it horizontal during 10-second trials with small continuous perturbations. After each trial, participants were asked whether they perceived the platform movement. Perturbation magnitudes were adjusted for the next trial based on their response. This process continued for each staircase until the termination criteria were met, at which point participants’ individual perception threshold was determined. Participants then performed ten 40-second trials on the stabilometer, two trials in each condition: without perturbation, perturbation at the 100%, 80%, and 50% of the individual’s perception threshold, and the pilot study’s minimum threshold. Balance performance was defined as time-in-balance ratio and RMS deviation angle from horizontal. Perception thresholds varied significantly between participants individuals, with an RMS acceleration ranging from 2.67 and 12.80 cm/s^2^. The results showed that perturbation magnitude has a significant correlation with variability in deviation angle (R=0.24, p=0.0038). The results suggest that some participants can perceive very small perturbations during a challenging balance task. Subthreshold perturbations, although very small, can influence balance performance.

**NEW & NOTEWORTHY:** To our knowledge, this is the first study to quantitatively measure the conscious perception threshold of platform perturbations while doing a balancing task. We found that young healthy adults with little similar prior experience in tasks close to stabilameter balancing task can detect small platform perturbations while doing such challenging task. The results also showed that platform perturbations below conscious threshold can influence the balance performance on the stabilometer.

## INTRODUCTION

Our ability to perceive balance perturbations plays an important role in how we control balance; when balance perturbations, or balance errors, are detected, rapid compensatory actions are taken to prevent falling (1–4). To respond effectively to balance perturbations, it is crucial to establish a sensory-motor framework for a typical range of errors and their similarity to the errors we usually experience(5,6). This framework enables the detection and prediction of deviations from this norm that could result in a loss of balance (7).

Balance control is achieved through the integration of multiple sensory systems, primarily proprioceptive, visual, and vestibular, which are weighted differently depending on the context and characteristics of perturbations (8–10). For instance, during static actions such as quiet standing or sitting, studies have consistently shown that proprioception and vision provide the most sensitive detection of external perturbation, whereas the vestibular system plays a secondary role (11,12). In contrast, under dynamic conditions, such as walking, proprioceptive sensitivity is reduced, reflecting lower neural excitability within spinal and corticospinal pathways compared to static actions (13–15). Moreover, the ability to perceive and respond to external perturbations is influenced not only by the sensory system involved but also by specific characteristics of the perturbation, such as displacement, magnitude, and the acceleration-deceleration profile of the perturbation (16,17). Collectively, these findings suggest that the nervous system modulates sensory contributions to balance in a context-dependent manner, optimizing postural control based on both the activity performed and the nature of the external disturbance.

Evaluating the ability to perceive sensory stimuli is a common method to diagnose impairment in sensory systems (e.g., vision, hearing). This is done using psychophysics methods, such as a staircase method, to determine perception threshold (18). A staircase method involves presenting stimuli of varying intensities to participants. When participants indicate that they can detect the stimulus, the intensity is decreased for subsequent trials; conversely, intensity increases if the participant is unable to detect the stimulus. This iterative process continues until a predetermined criterion is attained. A similar staircase method can be used to quantify people’s ability to perceive balance perturbations.

Previous studies identified conscious perception thresholds of sensory stimuli that contribute to detecting balance disturbances; however, they are limited because they mainly focus on everyday actions,(11,12,16,19) where the contribution of conscious attention is lower compared to when a new dynamic balance skill is required (20–22). In these studies, the conscious perception threshold has been found while people are usually skilled for such movements and need little explicit and conscious attention to handle external perturbations for maintaining balance. Here we argue that such experimental setup could be more suitable for scenarios where a new balance skill is required, which normally rely more on conscious attention to detect the external perturbations (20–22). In addition, previous studies have not investigated the influence of subthreshold perturbations on balance performance. Subliminal visual and auditory stimuli, below conscious threshold, may activate specific regions of the brain despite participants’ unawareness (23–25). However, it is not known if balance perturbations that are below the conscious perception threshold are still large enough to disturb balance. This knowledge may inform future studies on error-augmented practice, where participants are exposed to external perturbations that increase motor error without their awareness of perturbations.

The purpose of this study was to quantify the conscious perception threshold to external perturbations while performing a balancing task - standing on an unstable surface - in healthy young adults. Our secondary goal was to determine if ‘sub-threshold’ perturbations affect balance performance.

## MATERIALS AND METHODS

### Participants

Twenty-two healthy young adults (20-35 years) participated in this study. Participants were excluded if they had: difficulty understanding verbal or written English; or they reported any poor vision or poor hearing; any neurological or musculoskeletal condition; an injury that limits independent mobility; more than three months of training in dance, gymnastics, or other sporting or occupational activity with a large balance component; or participated in a moving platform study. The study was approved by the Research Ethics Board of the University Health Network (study ID: 21-6221), and participants provided written informed consent to participate. Participant age, sex, height, and mass were obtained.

### Procedure

All studies were completed within FallsLab in Toronto Rehabilitation Institute. The FallsLab platform is 6m x 3m and can move in all directions in the horizontal plane. The FallsLab platform was used to provide subtle, continuous balance perturbations. Participants’ balance performance was tested on a custom-made stabilometer platform that consisted of a swinging wooden platform (106 cm × 76 cm), which allows a maximum deviation of ± 30^°^ degrees to either side of the horizontal plane of the platform.

We designed 6 different platform movement waveforms where each waveform *s*^(*i*)^ is defined by a sum of two sinusoidal waves as follows:

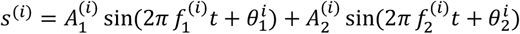

Where *A*_*1*_ and *A*_*2*_ are the amplitudes, *f*_*1*_ and *f*_*2*_ are the frequencies, and phases *θ*_*1*_ and *θ*_*2*_ are the phases of the two sine waves. We chose the first waveform based on a previous study (26). The parameters of the remaining 5 waveforms were then found such that the RMS of displacement, velocity, and acceleration of these waveforms were similar to those of the first waveform. Using a complex waveform that is the sum of two sinusoidal signals prevented participants from detecting and anticipating waveform patterns. The introduced balance perturbations were constructed by scaling these waveforms by a scaling factor *α*. Scaling the waveform by *α* increases or decreases the waveform magnitude, and results in scaling RMS of the waveform displacement, velocity, and acceleration with the same ratio across all waveforms. The balance perturbations were provided in the mediolateral direction. Waveform characteristics of all waveforms are shown in Table 1.

**Table 1.** Waveform characteristics at scaling factor of 1. A=sine wave amplitude; F=sine wave frequency; P=sine wave phase. Waveforms are chosen such that they have close RMS displacement, RMS velocity, and RMS acceleration

| Waveform | A<sub>1</sub><br>(cm) | F<sub>1</sub><br>(Hz) | P<sub>1</sub><br>(rad) | A<sub>2</sub><br>(cm) | F<sub>2</sub><br>(Hz) | P<sub>2</sub><br>(rad) | Peak<br>displacement<br>(cm) | Peak<br>velocity<br>(cm/s) | Peak acceleration<br>(cm/s<sup>2</sup>) |
| --- | --- | --- | --- | --- | --- | --- | --- | --- | --- |
| 0 | 2.5 | 0.250 | 0.11 | 3.0 | 0.500 | 3.22 | 4.77 | 9.90 | 34.33 |
| 1 | 2.6 | 0.246 | 1.99 | 3.2 | 0.491 | 3.99 | 5.18 | 13.89 | 34.91 |
| 2 | 2.3 | 0.235 | 13.94 | 3.5 | 0.470 | 3.86 | 4.19 | 13.26 | 35.41 |
| 3 | 2.0 | 0.236 | -9.99 | 3.4 | 0.471 | 9.74 | 5.40 | 12.06 | 30.25 |
| 4 | 2.2 | 0.237 | 1.08 | 3.4 | 0.475 | 3.77 | 3.69 | 12.46 | 35.16 |
| 5 | 2.4 | 0.257 | 0.33 | 2.8 | 0.514 | -6.58 | 5.11 | 12.52 | 31.24 |

Nine motion capture cameras (Vicon Motion Systems Ltd, Oxford, UK) surrounding the platform were used to track the positions and motion of reflective markers placed on the stabilometer and moving platform with a sampling frequency of 100 Hz. We attached 5 markers on the edges of stabilometer, 7 markers on the FallsLab platform, to track their movement and to compute the deviation angle of stabilometer during trials, and 5 markers to the participants’ head to track head movement. Video recordings were captured from 4 video cameras placed around the laboratory. These videos captured participants’ behavioural responses to the balance testing.

### Data collection

Participants initially completed one 30-second familiarization trial standing on the stabilometer mounted on the FallsLab platform without perturbations as they were naive to the task. Participants also performed one familiarization standing trial on the FallsLab platform with a large perturbation (α=1) to allow them to understand what the perturbations feel like. Participants then wore bottom-half masked goggles and noise-canceling headphones to minimize visual and auditory cues (e.g., from seeing the platform railing move relative to the surrounding walls). To ensure that participants did not tilt their heads to the rails of the platform, which provides additional visual cues for detecting perturbations, each participant performed a head calibration trial and tilted their head until they saw the rails. During data processing, we measured the minimum pitch angle using the data collected from this trial. If the participant had at least one second of head tilt lower than this minimum angle, they likely had seen the rails, and may therefore have received additional visual cues to indicate that the platform was moving during the trial.

During the test trials, participants stood on the stabilometer mounted on the moving platform and were asked to look forward at a focal point on the wall. Trials started when the stabilometer was in the horizontal position. There was a short pause between trials where the participant could step off the stabilometer. Longer rest breaks (∼1 min) were scheduled between blocks of trials (every 10-12 trials) to prevent fatigue. Additionally, participants were informed that they may request a rest break at any time.

To determine perception thresholds, participants experienced multiple 10-second trials, with or without platform movement. After each trial, participants were asked whether they perceived the platform movement. Perturbation magnitudes were scaled up in the next trial if participants could not perceive them and scaled down if participants could. The scale step size started from 0.4 and was divided by two after each reversal.

There were four perturbation waveforms (Waveforms 1-4; Table 2); two waveforms followed a descending staircase (starting from Scale 1) whereas two followed an ascending staircase (starting from Scale 0, i.e., no perturbation). For half of the participants Waveforms 1 and 3 were used for the ascending procedure and Waveforms 2 and 4 were used for the descending procedure, and vice versa for the other half of participants. Each block of five trials included one trial from each staircase and one catch trial with no perturbation, delivered in a pseudorandom order. The staircase ended when there were four reversals, and the perturbation perception threshold was defined as the average of the last two reversals. The participant’s individual perception threshold was selected as the minimum of the 4 thresholds from the 4 staircases. Figure 1 shows an example of the staircase procedure.

**Table 2.** Participant characteristics.

|  | Mean or count | Standard deviation | Minimum | Maximum |
| --- | --- | --- | --- | --- |
| Age (years) | 24.5 | 3.66 | 20.0 | 32.0 |
| Sex (number) |  |  |  |  |
| Men | 8 |  |  |  |
| Women | 9 |  |  |  |
| Height (cm) | 170.1 | 9.8 | 155.0 | 183.0 |
| Body mass (kg) | 74.3 | 23.5 | 49.5 | 109.5 |

**Figure 1.**
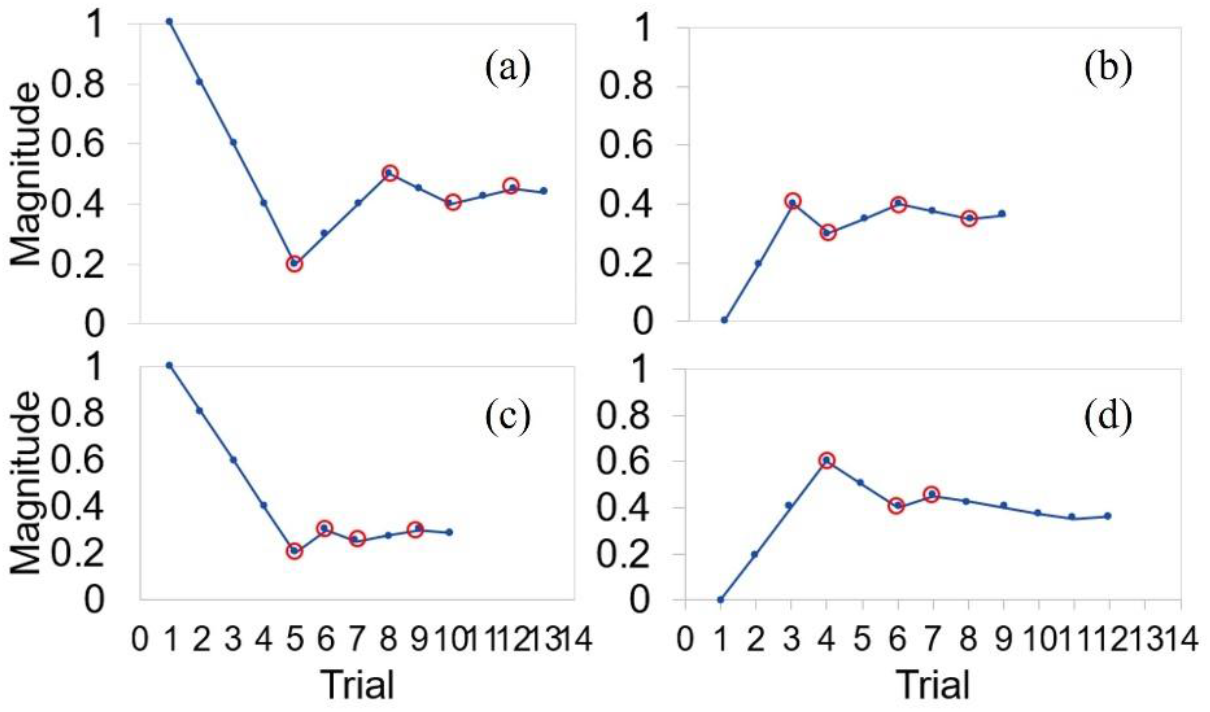
Staircase procedure. Multiple staircase procedures were performed for 4 different waveforms, 2 in descending (a and c) and 2 in ascending (b and d) orders. At each trial the magnitude increases or decreases based on the participant’s response. The procedure was terminated when four reversals happened (shown by red circles), and the threshold is computed as the average of the last 2 reversals.

Participants then performed 10, 40-second stabilometer trials, which included two trials (Waveforms 5, 6) in each of the following conditions: without perturbation, perturbation at 100%, 80%, and 50% of the individual’s perception threshold, and perturbation at the minimum perception threshold determined from a pilot study (*α*=0.16 from 6 participants) as a useful additional datapoint. The ten trials were presented in the same random order across all participants (50%, no perturbation, 80%, pilot, 50%, no perturbation, pilot, 100%, 80%, 100%).

The data collection session lasted approximately 2.5 to 3 hours, which allowed time for completing informed consent, instructing the participant, experimental trials, scheduled and requested rest breaks, setup and removal of equipment, and participant debriefing.

### Data processing

Kinematic data were filtered using a low-pass 4^th^ order zero phase lag Butterworth filter with a cutoff frequency of 10 Hz. Data were processed using MATLAB (2018a, The Mathworks Inc., Natick, Massachusetts, USA.)

We used two metrics to measure participant balance performance: 1) root mean square (RMS) of the deviation angle, and 2) time-in-balance ratio. RMS of deviation angle of stabilometer during each trial was computed as:

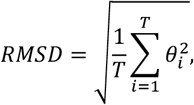

where *T* is the total trial time (in frames for a sampling rate of 100 frames per second) and *θ*_*i*_ is the deviation angle at frame *i*. The deviation angle in the frontal plane was computed from tracking markers attached to the stabilometer platform. Time-in-balance ratio was the amount of time that deviation angle was within ±2.5^°^ of horizontal, divided by the total trial duration. We excluded time in balance periods less than 20 frames (i.e., 200 ms) as these are considered as quick passes through horizontal, rather than periods where the stabilometer platform was held horizontal.

### Data analysis

Data analysis was conducted using SAS (Version 9.4, SAS Institute, Cary, North Carolina, USA). The dependent variables were balance performance measurements (RMS and time-in-balance), represented in two ways: 1) the original values of measurements including all perturbed and unperturbed (α=0) trials, and 2) the difference (delta) between balance performance in each perturbed trial and the average of performance of the participant in unperturbed trials. We used a generalized linear model to determine the relationship between the scaling factor and balance performance. We set the perturbation magnitudes corresponding to RMS acceleration of less than 0.07 cm/s^2^ to zero as there was no platform movement during these trials (confirmed from examining the platform acceleration signal derived from motion capture data).

Since all participants performed two trials with the same perturbation magnitude, we also compared balance performance in unperturbed trials (α=0) and perturbed trials with this magnitude (α=0.16) using a repeated measure ANOVA. Level of significance was 0.05 for all statistical tests.

## RESULTS

Four participants had a threshold with a root mean square (RMS) acceleration of less than 1 cm/s^2^. These participants had also a high false positive ratio (36% and more) in response to the catch trials, and therefore, were excluded from the study data analysis. Additionally, we excluded participants (n=1) who looked at the railing in >15% of trials. Participant characteristics for the remaining 17 participants are presented in Table 2.

For the remaining participants whose perception threshold was lower than the pilot threshold, the trials with pilot threshold magnitude were excluded from the analysis of balance performance in relation to perturbation magnitude (Objective 2), as our goal was to investigate the relationship between balance performance and perturbation magnitude below the perception threshold. We asked participants after each trial whether they felt the perturbations. The average true negative ratio among participants was 0.57.

### Perception thresholds

The mean±standard deviation of the threshold in scale α was 0.28±0.14 with a range from 0.12 to 0.60. This corresponds to an RMS acceleration of 6.06±2.94 cm/s^2^ with a range from 2.67 to 12.80 cm/s^2^. Descriptive statistics of threshold in scaling factor and RMS and peak of displacement, velocity, and acceleration are presented in Table 3.

**Table 3.**
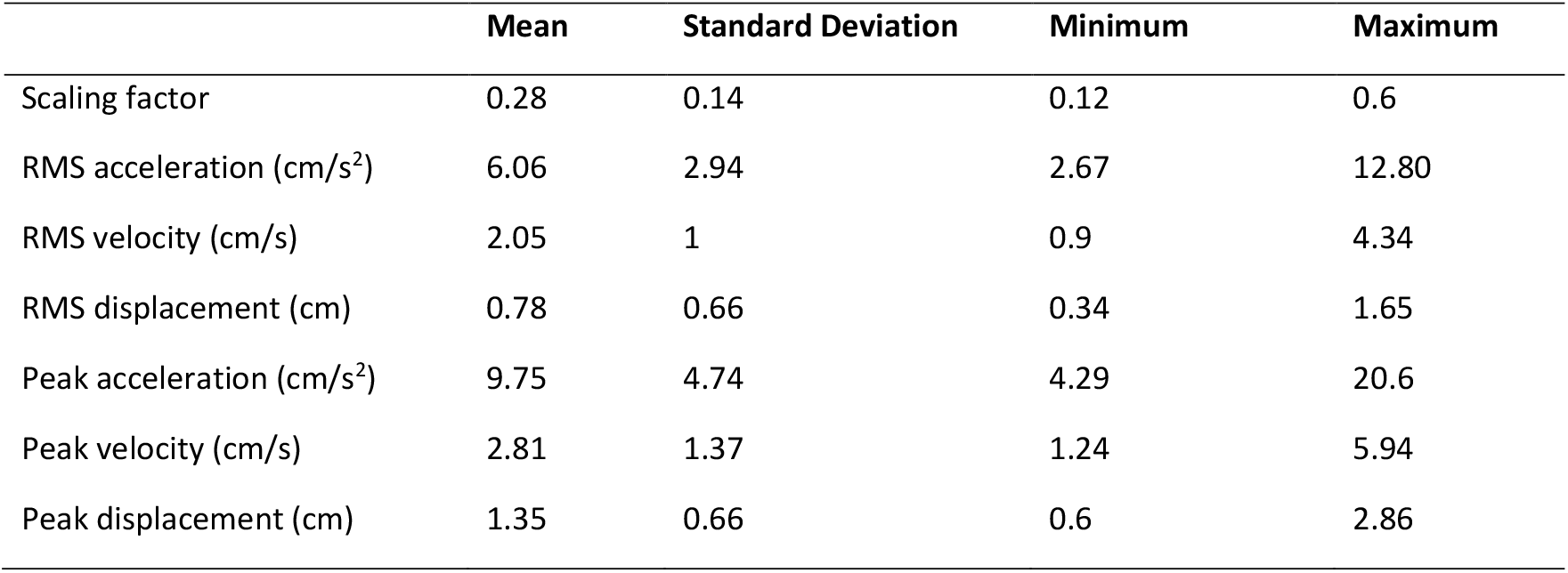
Descriptive statistics of thresholds in terms of scale and peak and rms displacement, velocity, and acceleration.

### Balance performance in relation to perturbation magnitude

The results showed no statistically significant correlation between time-in-balance ratio and perturbation magnitude (R=-0.18, p = 0.1195). For every one-unit increase in scale, there was an 11% decrease in the time-in-balance ratio (i.e., 4.4 seconds in a 40-second trial; Table 4). However, RMS deviation angle showed a statistically significant positive correlation with perturbation magnitude (R=0.24; p =0.0038), where for every one-unit increase in scale, there is an increase in RMS deviation angle of 2.57 degrees (Table 4).

**Table 4.** Generalized linear model which reflects the relationship between perturbation magnitude and balance performance in terms of time-in-balance ratio, delta time-in-balance ratio, RMS deviation angle, and delta RMS deviation angle. Values for the estimate include the 95% confidence interval in brackets. Z statistics indicate the standardized test values under the null hypothesis; p-values (Pr > |Z|) reflect statistical significance (p < 0.05 threshold).

| Variable | Estimate | Z | Pr > \|Z\| |
| --- | --- | --- | --- |
| **Time in balance ration** | -0.11 [-0.26, 0.03] | -1.56 | 0.12 |
| **RMDS (degrees)** | 2.57 [0.83, 4.31] | 2.89 | 0.0038 |
| **Delta time in balance ratio** | -0.042 [-0.21, 0.13] | -0.50 | 0.62 |
| **Delta RMSD (degrees)** | 3.4 [0.40, 6.4] | 2.22 | 0.026 |

There was no statistically significant correlation between delta time-in-balance-ratio and perturbation magnitude (slope=-0.042; p = 0.62). However, there was a statistically significant correlation between delta RMS deviation angle and the perturbation magnitude (slope=3.4; p = 0.026; Table 4).

There was a significant increase in time-in-balance ratio from balance performance perturbed with the magnitude informed from pilot study (α = 0.16) to unperturbed balance performance (F_1,15_ =10.39, p = 0.0053). Similarly, there was a significant decrease in RMS when comparing balance performance perturbed with the magnitude informed from pilot study (α = 0.16) and unperturbed balance performance (F_1,15_ =13.57, p = 0.002; Figure 2).

**Figure 2.**
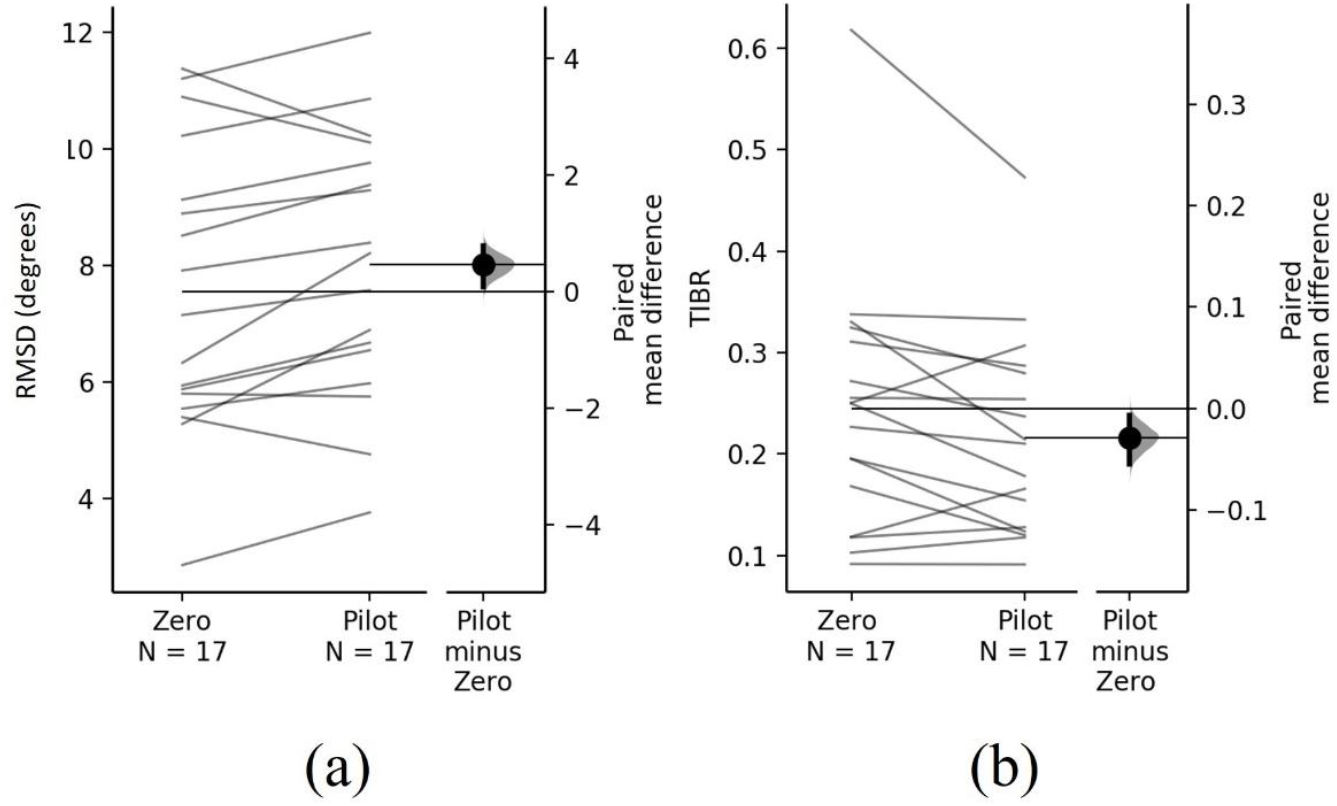
Effect of pilot threshold perturbations on balance performance. Comparing balance performance between the trials without perturbations and the trials where the participant was perturbed with a magnitude computed from the pilot study. Balance performance is measured using two metrics: a) RMS of deviation angle in degrees (RMSD) b) time-in-balance ratio (TIBR).

## DISCUSSION

The aims of this study were to quantify the conscious perception threshold of platform perturbation in healthy young adults while performing a stabilometer balance task, and to determine the effect of subthreshold perturbations on the balance performance. We found that this population can perceive subtle mediolateral perturbations while performing a novel and challenging balancing task with an average threshold of RMS acceleration 6.06±2.94 cm/s^2^.

Several studies have reported the perception threshold in the context of platform perturbations during quiet standing or sitting, with peak accelerations ranging from 0.8 to 2 cm/s^2^ (11,12,16,19). In our study, the peak acceleration at the perception threshold was 9.75 ± 4.74 cm/s^2^. Although many factors influence this threshold, such as waveform characteristics, trial length, and perturbation direction, the reported thresholds are significantly lower than those found in our study. Apart from the aforementioned differences, a primary distinction of our study is that, unlike quiet standing or sitting, the movement of the body and foot base due to platform perturbations is combined with roll rotation of the stabilometer surface. This combination makes it more challenging to detect these perturbations. Hlavaka et al. and Fitzpatrick et al. have suggested that proprioception and vision sensory systems contribute most significantly to detecting perturbations (11,12). In our study, we minimized the contribution of vision by asking participants to wear goggles that limited the lower half of the visual field. Also, previous studies indicate that, in a healthy population, disturbances detected from the foot base are primarily identified through proprioceptive feedback from the lower extremity (27,28).

However, in our study, keeping body upright while performing the stabilometer task enforces a large amount of effort from lower extremity musculature, potentially decreasing their contribution in perceiving movement caused by platform perturbations.

The vestibular system is reported to have the least contribution in perceiving balance perturbations (11,12). This is primarily because the acceleration of platform perturbations is not amplified from foot to head. Instead, this chain can be modeled as a multilink distribution, which dampens the force at both ends (16). Kingma showed that the threshold of the vestibular system for perceiving the direction of linear acceleration is approximately 6.5 cm/s^2^ for lateral movement and 8.5 cm/s^2^ for anterior-posterior movement during sitting (29). The similarity of the results of this study and ours could also suggest that in a stabilometer balance study where the visual and proprioceptive systems are limited, the main contributing sensory system is the vestibular system, resulting in a higher perception threshold compared the studies conducted on standing or sitting.

Preventing falls after a loss of balance requires a rapid reaction, involving unconscious response using local muscle reflexes or later brainstem responses, and even later conscious cortical responses (30). For populations with impaired spinal and brain-stem responses, the importance of conscious perception of external perturbations using cortically mediated and prefrontal responses becomes more significant (1,2,31). There are also several studies that corroborate this and suggest that populations such as older adults (32,33), and people with Parkinson’s disease (34,35), or stroke (36), have a degraded ability to consciously perceive perturbations. We designed our study based on a novel balancing task, which requires more conscious effort for maintaining balance compared to everyday activities such as standing or walking. Therefore, our findings may primarily apply to novel balance tasks compared to studies that found similar thresholds for everyday activities, where balance is mainly maintained unconsciously.

To our knowledge, no previous studies investigated the influence of subthreshold perturbations on balance performance. Our results showed that, even in ranges below the conscious perception threshold, perturbation magnitude and RMS deviation angle have an inverse relationship showing an immediate impact of perturbations in this range on balance. This suggests that external perturbations below the conscious perception threshold may also need automatic corrective responses to stabilize posture. Unlike RMS deviation angle, our results did not show a significant correlation between perturbation magnitude and time-in-balance ratio. One explanation is that these perturbations may disrupt fine postural control, increasing the deviation angle inside the range of measured time-in-balance (±2.5^°^). Time-in-balance is often an all-or-nothing measure with ceiling effects, making it less sensitive to gradual changes in control. Thus, deviation angle better captures subtle performance degradation under increasing perturbations.

Apart from such immediate impact, subthreshold perturbations may also impose a longer-term influence on learning, future behaviour and balance strategies. These perturbations can be viewed as a subliminal stimuli, below conscious threshold, that impact on specific regions of the brain (23–25). These findings may also have implications for study of error-augmented practice; exposing participants to external perturbations that increase motor error without participants’ awareness of the perturbations may help us to understand the role of externally-applied versus internal errors in learning novel motor skills.

We excluded 4 participants with a very low threshold and a high false positive ratio in catch trial responses. Such scenarios could be one of the disadvantages of staircase method where the results are less robust to response biases compared to two-alternative-forced-choice (2AFC) method (18). Although 2AFC explicitly minimizes response bias, it typically needs a greater number of trials compared to adaptive staircase methods. In our study, we needed to take practice effects and fatigue into consideration, which made using 2AFC infeasible. In contrast, adaptive staircase methods achieve greater efficiency by concentrating trials near the threshold region where sensitivity transitions occur most sharply. Selectively refining the stimulus around this critical boundary minimizes unnecessary trials at extremes of the stimulus range that are either too detectable or imperceptible. In adaptive staircase methods, there is a trade-off between accuracy and required number of trials of the procedure based on the configuration parameters. In our study we set these parameters such there was a balance between these two. For example, large step sizes reduce the number of trials but can overshoot the threshold, decreasing accuracy, while small step sizes improve precision but require more trials. We also set number of reversals to 4. More number of reversals could increase accuracy but with the cost of more number of trials. In our study, the total staircase procedure (for all 4 staircase procedures for 4 waveforms) for participants was performed in an average of 46 trials (including catch trials) which took approximately 30 minutes. Each participant had 30 seconds rest time after each trial, and 1-2 minutes rest after each block of 10 to minimize the effect of physical and cognitive fatigue.

## DATA AVAILABILITY

Source data for this study are not publicly available due to privacy or ethical restrictions.

## AUTHOR CONTRIBUTIONS

K.M., L.T., A.N., and A.M conceived and designed research; K.M. analyzed data; K.M. and A.M. performed experiments; K.M. interpreted results of experiments; K.M. prepared figures; K.M., L.T., A.N., and A.M. drafted manuscript; K.M., L.T., A.N., and A.M. edited and revised the manuscript; K.M., L.T., A.N., and A.M. approved final version of the manuscript;

